# FRET-based screening in HEK293T identifies p38 MAPK and PKC inhibition as therapeutic targets for α-synuclein aggregation

**DOI:** 10.1101/2021.02.16.431382

**Authors:** Alexander Svanbergsson, Fredrik Ek, Isak Martinsson, Jordi Rodo, Di Liu, Edoardo Brandi, Caroline Haikal, Laura Torres-Garcia, Wen Li, Gunnar Gouras, Roger Olsson, Tomas Björklund, Jia-Yi Li

## Abstract

Aggregation of α-synuclein is associated with neurodegeneration and a hallmark pathology in synucleinopathies. These aggregates are thought to function as prion-like particles where the conformation of misfolded α-synuclein determines the induced pathology’s traits similar to prion diseases. Still, little is known about the molecular targets facilitating the conformation-specific biological effects, but their identification could form the basis for new therapeutic intervention. High-throughput screening (HTS) of annotated compound libraries could facilitate mechanistic investigation by identifying targets with impact on α-synuclein aggregation. To this end, we developed a FRET-based cellular reporter in HEK293T cells, with sensitivity down to 6.5 nM α-synuclein seeds. Using this model system, we identified GF109203X, SB202190, and SB203580 as inhibitors capable of preventing induction of α- synuclein aggregation via inhibition of p38 MAPK and PKC, respectively. Our findings highlight the value HTS brings to the mechanistic investigation of α-synuclein aggregation while simultaneously identifying novel therapeutic compounds.

## Introduction

Alpha-synuclein (α-syn) aggregation is a common feature of synucleinopathies, a group of neurodegenerative diseases including Parkinson’s Disease (PD), Multiple System Atrophy (MSA), and Dementia with Lewy Bodies (DLB). The deposition of α-syn aggregates presents distinctly depending on the disease. PD and DLB are both characterized by aggregates called Lewy Bodies (LB) and Lewy Neurites (LN) found in neurons (Parkkinen et al., 2008). MSA presents instead with predominantly glial cytoplasmic inclusions (GCI) in oligodendrocytes, but also a degree of neuronal cytoplasmic and intranuclear aggregates (Nishie et al., 2004).

Current hypotheses suggest synucleinopathies are driven by a cell-to-cell spread of misfolded α-syn in a prion-like manner. This mechanism is supported by a progressive temporal spread of synuclein pathology to interconnected brain areas following injection of fibrillar α-syn seeds into the striatum or olfactory bulb (Paumier et al., 2015; Rey et al., 2018). Pathology may also originate in the gastrointestinal tract following injection of recombinant fibrils or patient-derived aggregates and progress towards the brain, in line with Braak’s hypothesis (Braak et al., 2003; Holmqvist et al., 2014). The spread of pathology has also been reported in patients receiving grafts of fetal mesencephalic dopaminergic neurons, where Lewy pathology was observed during post-mortem autopsy 11-24 years following transplantation (Kordower, Chu, Hauser, Freeman, et al., 2008; Kordower, Chu, Hauser, Olanow, et al., 2008; J.-Y. Li et al., 2008; W. Li et al., 2016).

A prion-like spread may also account for the heterogeneity of synucleinopathies, as conformational differences may result in altered interactions. Like prion diseases, α-syn has been shown to take on various conformations upon misfolding, which in turn could alter the disease aetiology similar to bona fide prion diseases. Such structurally distinct fibrils have been produced in cell-free conditions and were shown to possess a structure-dependent spread of pathology and detrimental effects (Peelaerts et al., 2015). Differences in biological activity are also observed for aggregated α-syn isolated from PD and MSA patient brains, where MSA-derived aggregates display a higher seeding efficacy (Prusiner et al., 2015). These conformational and function differences of such pathological aggregates may form as a consequence of the local environment of the affected cell types, as recently reported (Peng et al., 2018).

While our understanding of the mechanisms involved in α-syn related pathology has vastly expanded over the last decade, this has not yet translated into treatment options for any synucleinopathy. Immunotherapy (Henderson et al., 2020), gene therapy (Heiss et al., 2019), or anti-sense oligonucleotides (Mercuri et al., 2018) are exciting treatment options for neurodegenerative diseases currently under development. Alternative approaches such as small molecule compound screening have also helped identify novel targets or treatments (Halliday et al., 2017). Annotated libraries with thousands of compounds have been established and are commercially available for the application for appropriate disease models.

To provide a high-throughput screening (HTS) platform, we have leveraged a system based on fluorescence resonance energy transfer (FRET)-based detection of α-syn aggregation by flow cytometry. After validation and characterisation of our method, we screened a library of small molecule kinase inhibitors. We identified three novel compounds with strong inhibitory effects on seeded α-syn aggregation *in vitro* by targeting p38 mitogen-activated protein kinase (p38 MAPK) and protein kinase C (PKC). We further examine the cellular alterations induced by the inhibitors and observe lysosomal-related changes are likely the source of the inhibitors’ protective effects.

## Results

### Generation and characterization of a FRET-based reporter system for α-syn aggregation

To generate a FRET-based α-syn aggregation reporter, we transduced HEK293T cells with lentiviruses encoding α-syn^A53T^-CFP and α-syn^A53T^-YFP. We selected the A53T mutant variant of α-syn, partly due to its relevance as one of the autosomal dominant variants of PD (Langston et al., 1998), but also due to the increased propensity to aggregate (Cabin et al., 2005; Kang et al., 2011). By fluorescence activated cell sorting (FACS) sorting on the double-positive cell population, we generated 12 monoclonal cell lines. The monoclonal lines, established by single-cell sorting and subsequent clonal expansion, were treated with PFFs and liposomes to facilitate direct delivery of aggregates and induce aggregation (Figure 1A). The final reporter cell line was selected based on normalized FRET intensity (FRET mean fluorescence intensity multiplied by %FRET positive) and signal-to-noise ratio of the FRET signal (data not shown) as detected by flow cytometry. The cell line is from here-on referred to as NPR-H-001.

**Figure 1.**
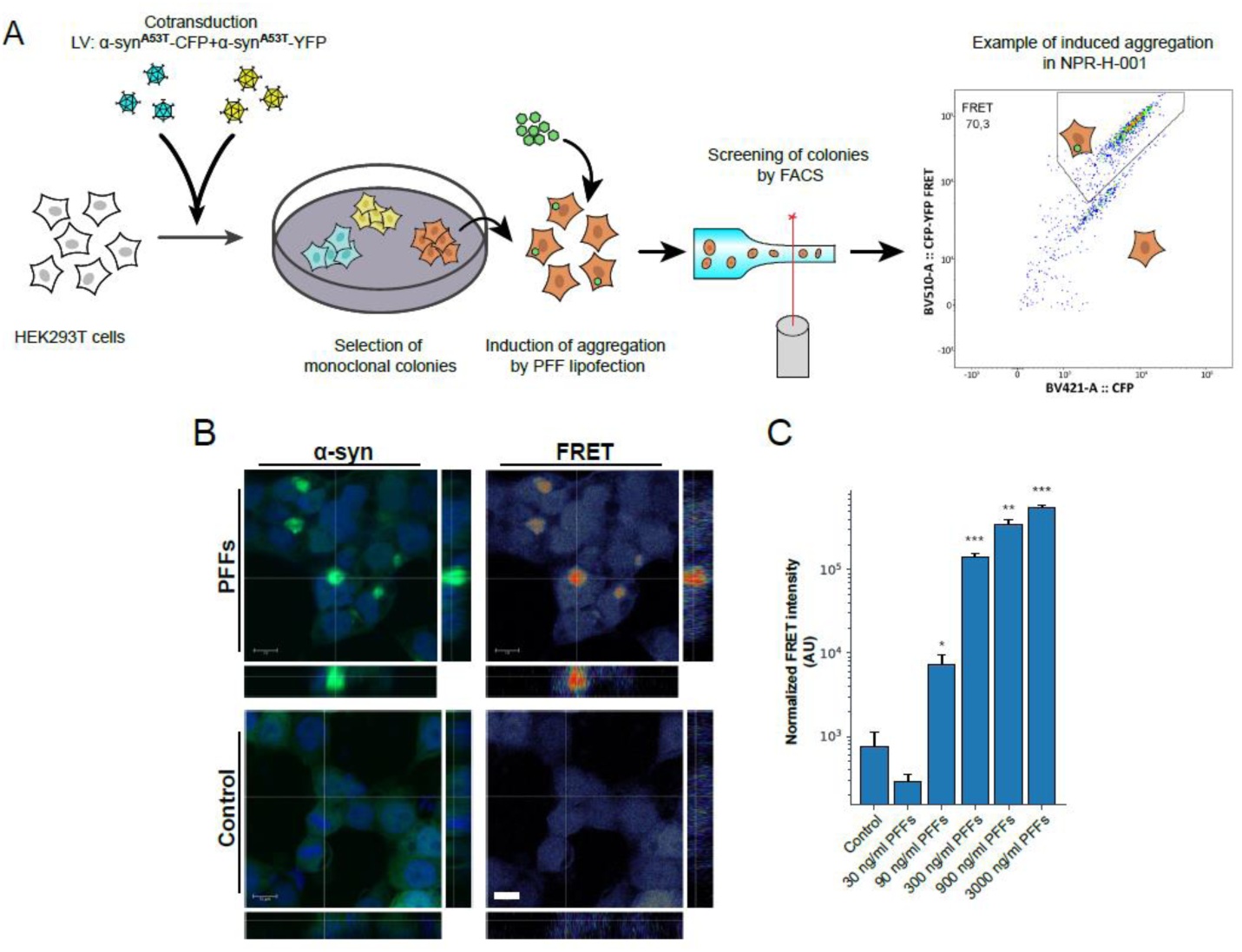
Monoclonal α-syn aggregate FRET reporter cells allow inclusion-specific detection of aggregates. (A) HEK293T cells were simultaneously transduced with lentivirus encoding α-synA53T-CFP and –YFP. Monoclonal cell lines were established by FACS single-cell sorting. Clonal lines were compared for brightness and signal-to-noise ratio of FRET by induction of aggregation by lipofection of α-syn PFFs to the biosensor lines.(B) Confocal microscopy confirmed a strong induction of FRET signal overlapping with the YFP signal, and no induced FRET signatures when treated with monomeric α-syn. (C) Sensitivity assessment by PFFs titration and FRET detection by flow cytometry (n=3). Bar chart values show mean ± SD, *p<0.05, **p<0.005, ***p<0.001. Statistical testing was performed using Brown-Forsythe and Welch one-way ANOVA for multiple comparisons to a control group.

As flow cytometry detects FRET signal on a per-cell basis, we sought to confirm the observed FRET signal originated from induced aggregates. NPR-H-001 cells were treated with PFFs and liposomes, followed by fixation 24 hours after addition for confocal microscopic visualisation (Figure 1B). The detected FRET signal was confirmed to mainly derive from the induced aggregates. We next sought to determine the sensitivity by which induced aggregation could be detected in NPR-H-001 reporter cells. Inducing aggregation with increasing amounts of PFFs delivered by liposomes, we significantly detect induced aggregation by PFFs by concentrations as low as 90 ng/ml (p=0.039). We further assessed the z-factor of our NPR-H-001 reporter cells, a parameter frequently used to evaluate suitability to HTS, reflecting the separation between the control and maximal induction. Using NPH-H-001 we obtained a z-factor of 0.79 when induced with 3000 ng/ml PFFs, indicating the assay is well suited for screening purposes.

### Induced aggregates recapture hallmarks of synucleinopathies

In the reporter cell line generation and characterization, we have applied liposomes for the delivery of PFFs. Liposome-mediated delivery facilitates the cytosolic entry of PFFs prior to initiation of aggregation, precluding the cytosolic entry from being studied. Therefore, we sought to assess if the direct addition of PFFs to the media could offer a comparable induction of aggregation, including endocytic events in the experimental paradigm. To this end, we compared the capacity of the two methods of inductions to induce α-syn seeding and deposition of insoluble α-syn.

As lipofection seeds aggregation more rapidly, cells treated with liposomes and PFFs were harvested 24 hours after addition compared to direct addition after 48 hours. We then performed sequential extraction of the soluble and insoluble fractions from protein lysates of HEK293T cells with induced α-syn aggregation or control, and assessed differences in accumulation of insoluble and phosphorylated α-syn (p-α-syn) via direct addition or lipofection-mediated delivery (Figure 2A). While the fraction of triton-X100 soluble p-α-syn remained unaltered, a 3.7-fold and 4.4-fold increase was observed in the insoluble fraction of direct addition and lipid delivery, respectively (Figure 2A). The formation of insoluble aggregates seen for both delivery methods suggests that both are viable options for studying α-syn aggregation using our novel NPR-H-001 reporter cells. Therefore, we used direct addition for all experiments to induce aggregation to allow uptake and internalization effects to be reflected in our model.

**Figure 2.**
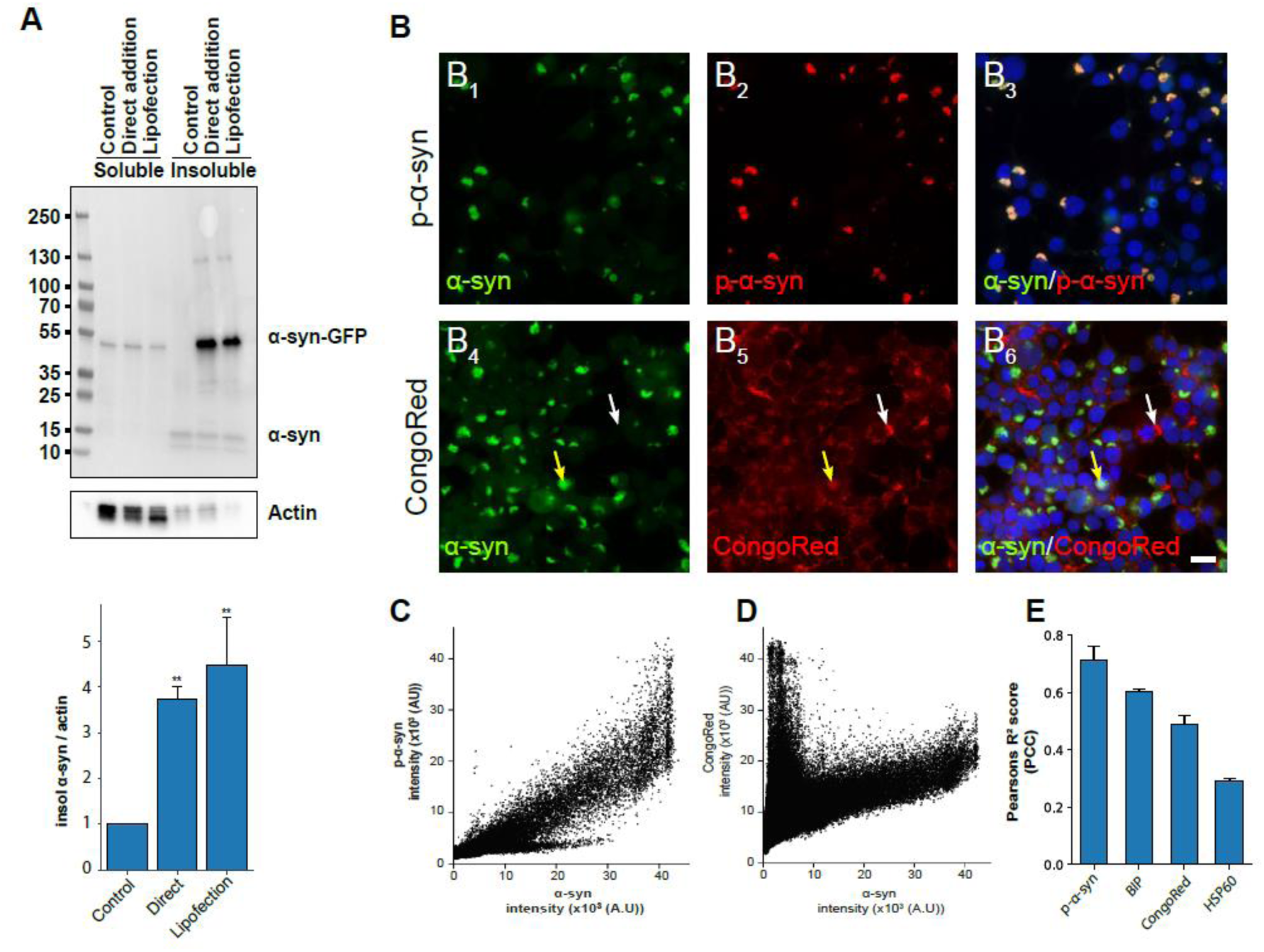
Induction of α-syn aggregation in NPR-H-001 cells recaptures key hallmarks of synucleinopathy. (A) Treatment of Hek293TA53T-GFP with PFFs leads to a significant increase in phosphorylated α-syn compared to treatment with monomeric α-syn. Quantification of Western blot analysis showed levels of phosphorylated α-synuclein increased 3.7 (p<0.009) and 4.4 (p<0.003) folds with direct addition and lipofection, respectively, when compared to control treatment (n=3). (B) α-Syn forms aggregates visible as compacted green inclusions, and colocalizes with phosphorylated α-synuclein as seen in B3. Staining with CongoRed reveals two populations of aggregates, double-positive for GFP and CongoRed (B6 Yellow arrow) and CongoRed positive GFP negative (B6 White arrow) (Scale bar: 20 µm). (C-D) Correlation scatter plot between α-syn-GFP and p-α-syn (C) and CongoRed (D). (E) PCC (R2) for p-α-syn (0.71±0.05), CongoRed (0.48±0.03) and the chaperone proteins BIP (0.60±0.01) and HSP60 (R2=0.29±0.008) (n=3). Bar charts show mean ± SD, *p<0.05, **p<0.005, ***p<0.001. Statistical testing was performed using one-way ANOVA with Dunnett’s T3 post hoc test for multiple comparisons to a control group, except (E) where comparisons between groups were not meaningful.

We further characterized the induced aggregates for markers of pathology related to synucleinopathy. First, we assessed the extent to which induced aggregates were phosphorylated by immunostaining with anti-phosphor (S129)-α-syn (pS129). We observed a strong positive correlation defined by Pearson’s correlation coefficient (PCC) (R^2^=0.71±0.05) (Figure 2 B^1-3^ and 2C). Aggregated α-syn is also known to be rich in cross-β-sheet structure as this is the major fold of amyloids. Staining for the amyloid structure with CongoRed resulted in a moderate overall colocalization (R^2^=0.48±0.03). However, two populations of CongoRed staining were observed, as evident by the correlation scatterplot (Figure 2B^4-6^ and 2D). One population with intense CongoRed staining and no GFP intensity (Figure 2B^4-6^ white arrow) and the other with moderate CongoRed but high GFP intensity (Figure 2B^4-6^ yellow arrow). These observed populations likely stem from extracellular PFFs which still have not been internalized and intracellular seeded aggregates. Additionally, we also stained for BIP and HSP60 as these are chaperones frequently associated with protein misfolding. We observed a moderate correlation of the induced aggregates with the chaperones BIP (R^2^=0.60±0.01) and HSP60 (R^2^=0.29±0.008), respectively (Supplemental Figure 1). Taken together, the analyses of the induced aggregates indicate that key pathological features are shared with aggregates found in synucleinopathies, proving the relevance of this seeded aggregation model as an experimental tool for studying synucleinopathies.

### High-throughput screening identifies potential modulators of α-syn aggregation

To identify possible modulators of α-syn aggregation and showcase the NPR-H-001 model for HTS, we performed a small molecule compound screening. We selected a library of kinase inhibitors for kinases that previously have been implicated in PD pathology (Esteves & Cardoso, 2017).

NPR-H-001 reporter cells were pre-treated for 30 min. with 10 µM inhibitors of the compound library prior to the addition of PFFs. At the 48-hour endpoint following addition of PFFs, cells were fixed and assessed for FRET signal by flow cytometric analysis (Figure 3A). We calculated the z-Factor across all three experimental repetitions, resulting in a z-Factor of 0.48±0.1 (mean±SD), suggesting that the assay is well suited for HTS.

**Figure 3.**
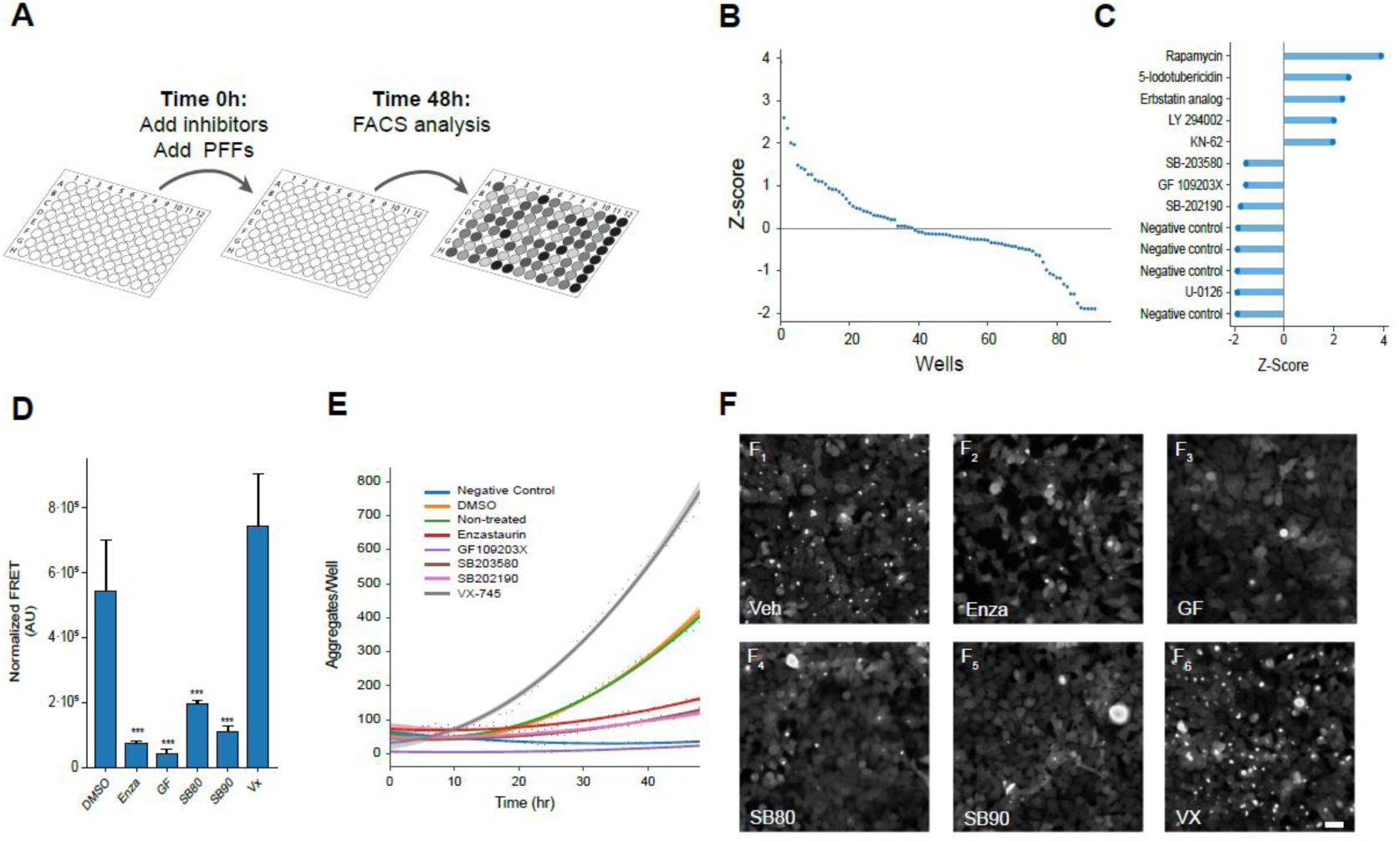
High-throughput screening identifies three kinase inhibitors with potent preventative effects on α-synuclein seeding. (A) Schematic workflow of kinase inhibitor screening using NPR-H001 cells. Assay plates seeded with reporter cells are pre-treated with inhibitors prior to PFFs addition and analysed by flow cytometry after 48 hours. (B) Resulting z-scores from kinase inhibitor screening distributes as a waterfall plot with the majority displaying little effect and few tail-end candidates showing large alterations. (C) Ranked representation of all samples with z-scores beyond the 1.5 threshold identifies compounds with potential impact on α-synuclein aggregation. (D-F) Single compound validations show a visual reduction in formed α-syn aggregates 48 hours following PFFs addition (scale 40 µm) (D), with differences already being detectable 20h post PFFs addition (E). (F) Using NPR-H-001 reporter cells to confirm the significant reduction in induced aggregation (Enza: p<0.001, GF: p<0.001, SB80: p<0.001, SB90: p<0.001) following compound addition by flow cytometry. Bar chart show mean ± SD, ***p<0.001. Statistical testing was performed using one-way ANOVA with Dunnett’s T3 post hoc test for multiple comparisons to a control group.

The success criteria for hit-compounds were defined as a z-score of -1.5, with negative controls obtaining an average z-score of -1.88±0.01 (mean±SD) indicating hit compounds will have a strong preventative effect on induced α-syn aggregation (Supplementary Table 1). Beyond the threshold, we identified 4 kinase inhibitors (SB203580 (SB80), GF109203X (GF), SB202190 (SB90) and U-0126) with strong reduction of normalized FRET signal. Of the four candidate compounds, two were inhibitors of p38 MAPK, one inhibits PKC, and the last targets MEK. We also observed 5 compounds with z-scores surpassing vehicle control (rapamycin, 5-iodotubericidin, Erbstatin analogue, LY29002 & KN-62), indicating an exacerbation of the aggregate phenotype (Figure 3B-C).

To validate the aggregation-inhibition potential of candidate compounds, we assessed the efficacy for each inhibitor by flow cytometry and live-cell microscopy. For comparison, we included two additional inhibitors that have previously been used in clinical testing, though not for in relation to α-syn, with the same targets as the identified compounds, Enzastaurin targeting PKC (trial id: NCT03263026) and VX-745 (VX) targeting p38 MAPK (trial id: NCT02423200)

First, we treated NPR-H-001 cells with inhibitors and assessed the aggregation state by flow cytometry, in the same way as was performed for the screen. The compound treatment resulted in a significant decrease in aggregate load for Enza, GF, SB80 and SB90 (Enza 7.4 fold, p<0.001; GF 12.4 fold, p<0.001; SB80 2.7 fold, p<0.001; SB90 4.8 fold, p<0.001). Contrary to the other inhibitors, VX displayed no significant change in aggregate load (Figure 3D).

We next sought to assess the impact of inhibitor treatment of the kinetics of induced α-syn aggregation. For this, we performed live-cell microscopy with a HEK293T cell line with the single α-syn^A53T^-GFP fusion construct. Similar to the assessment by flow cytometry, a marked reduction in the quantity of α-syn aggregates was observed by live-cell microscopy for all inhibitors tested except VX (Figure 3E-F). Rather VX led to a significant increase in aggregate load and early detection (∼10 hours following PFFs addition) (Figure 3E).

Taken together, the outcome of the kinase inhibitor screen shows the applicability of the NPR-H-001 reporter for HTS, where we identified 3 compounds with strong inhibition of α-syn aggregation by targeting p38 MAPK (SB80 and SB90) and PKC (GF).

### Changes in the phosphorylation state of endocytosis-related proteins differentiate PKC from p38MAPK inhibitors

To investigate the changes induced by the kinase inhibitors, we compared the phosphor-proteome of inhibitor-treated cells against vehicle-treated controls. As all identified inhibitors were targeting kinases, immediate effects of compound addition would be expected in an altered phosphorylation profile. To this end, we collected lysate from inhibitor-treated cells and enriched part of the lysate for phosphoproteins. The enriched protein sample was analysed by mass spectroscopy to identify differentially phosphorylated proteins.

We detected 11620 phospho-epitopes from a total of 4035 proteins, with 3230 epitopes identified as differentially regulated in the inhibitor-treated cells compared to the control. In line with our expectations, clustering of samples based on the treatment is observed (Figure 4A). The high correlation (R^2^<0.9) among replicates for each inhibitor supports the reproducibility of the treatment. However, we also observe elevated correlation between inhibitors with the same target indicating similarity in the altered phosphor-proteome (Figure 4B). The target specificity is also reflected in the overlap of significantly altered phospho-epitopes with a minimum log-fold change (LFC) of 0.5, where 44.5% and 60.8% are shared between the p38 and PKC targeting inhibitors, respectively (Figure 4C).

**Figure 4.**
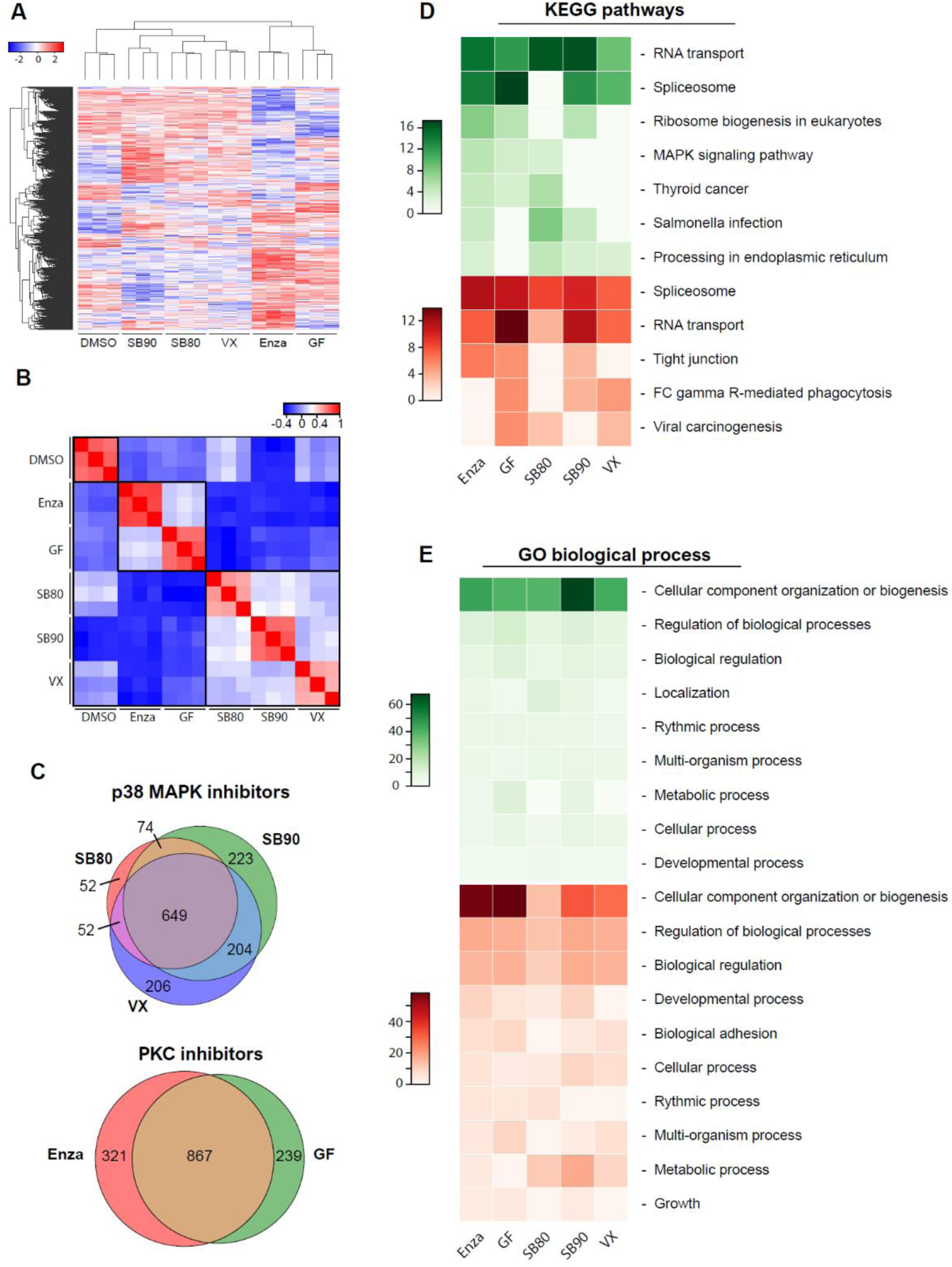
**Phospho-proteome shows alterations in protein translation and cellular component organization/biogenesis following inhibitor treatment**. (A) Unsupervised clustering of the altered phospho-proteome groups samples correctly based on target pathways. (B) Reproducibility between replicates and similarity among inhibitors with the same targets give rise to defined clusters based on PCC. (C) Contrasting the altered phospho-proteome depending on inhibitor target shows a high degree of overlap, with 44.5% for p38 MAPK and 60.8% for PKC. (D-E) Functional analysis using DAVID of the altered phospho-proteome identified two main processes depending on the analysis employed. (D) KEGG analysis indicated the main alterations, both for increased- and decreased phosphorylation being RNA transport and spliceosome. In contrast, GO analysis displayed only one major profile found in both negatively and positively altered, being cellular component organization and biogenesis.

Using the altered phospho-proteome we performed functional analysis to identify altered molecular pathways and processes using KEGG pathway analysis and Gene Ontology (GO) biological processes, respectively (Figure 4D-E). From KEGG pathway analysis, the strongest profile altered for all compounds is related to transcription/translation. Both spliceosome and RNA transport were among the most altered, both negatively and positively regulated (Figure 4D). Interestingly, the PKC targeting compounds also lead to increased MAPK signalling, as well as for SB80 (Figure 4D). Analysis of altered biological processes with GO identified cellular organization and biogenesis, as the main process with altered phosphorylation. These results indicate that translation machinery and organelle biogenesis are the major altered pathways in response to the treatments with the inhibitory compounds.

### Proteome analyses indicate alterations in endo-lysosome pathways upon PKC and p38 MAPK inhibition

With protein production and the translation machinery mostly affected within the phosphor-proteome, we sought to investigate changes in the global proteome.

From the global proteome, we detected 4989 proteins across all samples, with 1183 proteins being significantly altered when compared to untreated samples. Similar to the phospho-proteome, unsupervised clustering results in grouped inhibitors based on treatment and inhibitor target (Figure5A), with the exception of SB90. SB90 branches off from all other inhibitors but does however show elevated PCC with SB80 and VX (Figure 5B). The overlap between the altered proteome for each inhibitor target drops compared to the phospho-proteome, as the proteome only shares 27.7% and 46.4% overlap of significantly altered proteins for p38 MAPK- and PKC-inhibition, respectively (Figure 5C). This represents a 16.8% and 14.4% decrease in the overlap, respectively.

**Figure 5.**
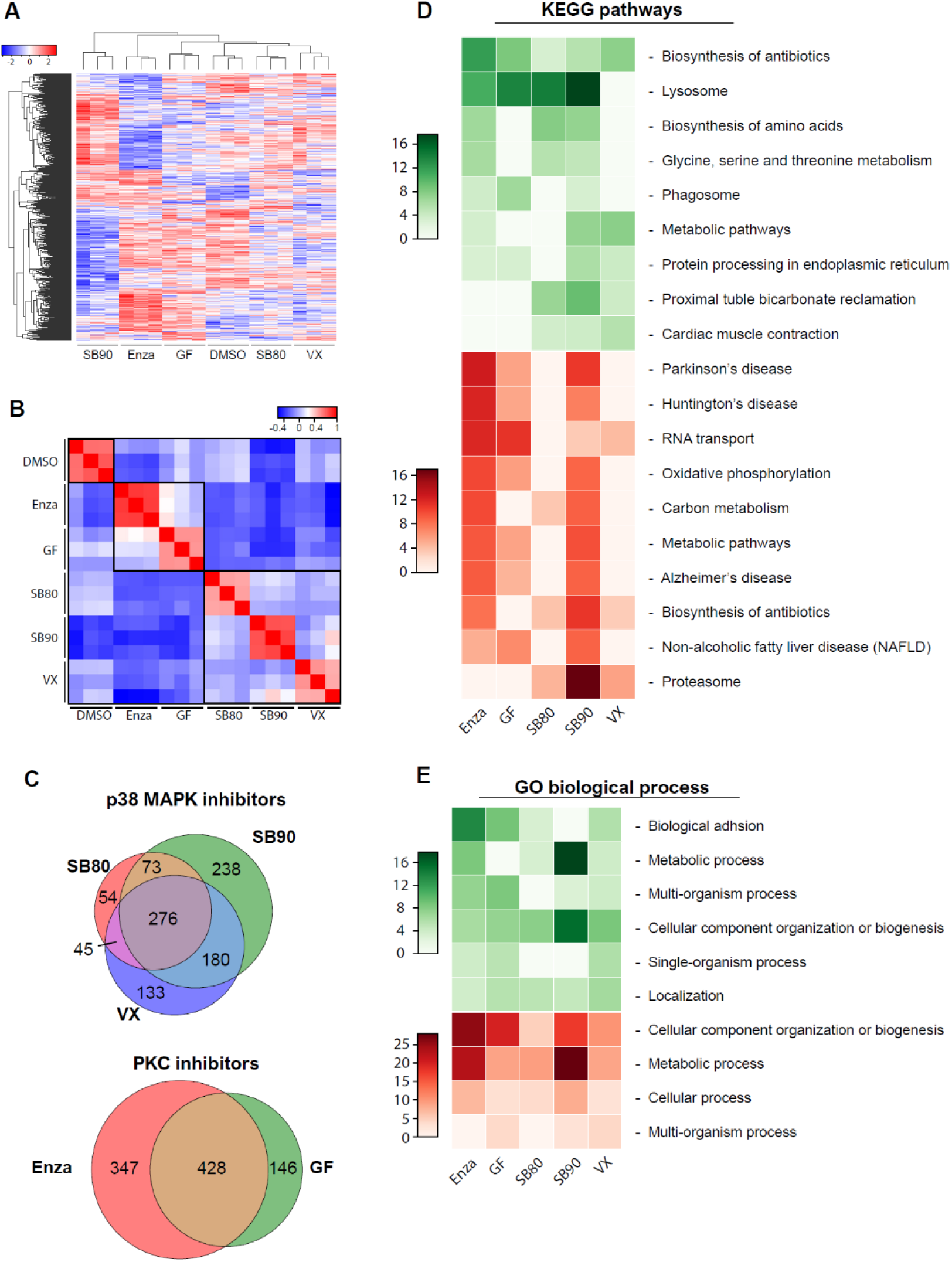
Proteome analyses indicate altered protein levels of lysosome pathways in protective phenotype. (A) Unsupervised clustering of the significantly altered proteome groups clusters correctly based on inhibitor and inhibitor target except for SB202190, which is grouped alone. (B) High sample correlation within inhibitor treatment confirms reproducibility, while clusters are also observed within inhibitors with the same molecular target. (C) The overlap among the significantly altered proteins of samples treated with inhibitors of the same target is 27.7 % and 46.4 % for p38 MAPK and PKC, respectively, indicating dissimilar effects of the inhibitors. (D) KEGG pathway analysis reveals downregulation of protein levels within neurodegeneration-related KEGG pathways for 3 of 5 compounds. However, lysosome-related pathways show enrichment for 4 of 4 inhibitors with protective phenotype. (E) GO: Biological processes remain devoid of alterations matching the observed protective phenotype.

To identify pathways or molecular events which may be involved in the observed protective phenotype, we performed functional analysis based on the altered proteome. Signatures of particular interest would have to be present for all inhibitors except VX, which did not display the protection from induced α-syn pathology. KEGG pathway analysis, as seen in Figure 5D, displayed an increase in proteins related to the lysosomal pathway for all compounds with preventative effects on induced α-syn aggregation. This suggests a further investigation of lysosome-related pathways may be warranted.

Of further interest is the decrease abundance of proteins in the pathways mapped in the KEGG database for PD, Alzheimer’s Disease (AD), and Huntington’s Disease (HD). On further examination of the significantly depleted proteins mapping to these pathways, most were found to relate to mitochondrial-related function and metabolism.

### PKC and p38 MAPK inhibition induces alterations in endo-lysosomal compartments

To dissect the detailed machinery altered in the endo-lysosomal system as indicated by the altered proteome, we quantified the abundance of acidified compartments. We first visualized acidified compartments by staining inhibitor-treated HEK293T cells with lysotracker, a compound stain with pH-dependent fluorescence intensity. The abundance of acidified compartments was visualized by live-cell microscopy and quantified by flow cytometry. Significant elevation in acidified compartments was observed for both PKC targeting inhibitors as well as for SB90 targeting p38 MAPK, although to a lesser extent (Figure 6A-B).

**Figure 6.**
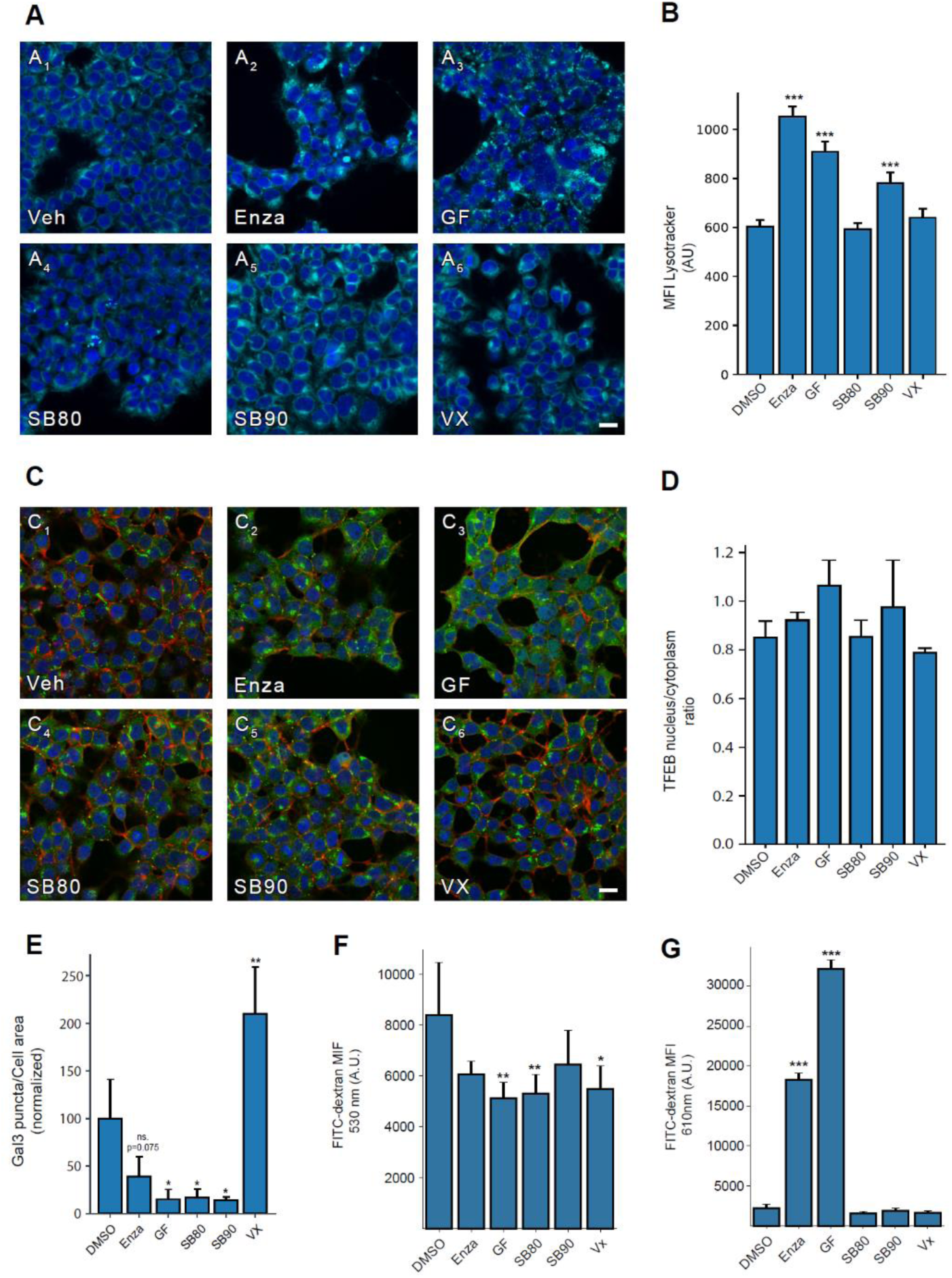
Lysosome and intracellular vesicular alterations in cells treated with hit-compounds. (A) Visualization of acidified compartments by lysotracker reveals compartments with increased brightness and a more puncta-like appearance. (Scale 20 µm) (B) The changes in acidified compartments are found to be statistically significant upon flow cytometric analysis for Enza (p<0.001), GF202190 (p=0.001), and SB90 (p=0.001). (C-D) Analysis of nuclear translocation at TFEB at 48h following inhibitor treatment reveal no significant changes in the ratio of nuclear TFEB to cytoplasmic TFEB. (Scale 20 µm) (E) Integrity of the endo-lysosomal compartments was examined by galectin3-GFP spot formation 48 hours following PFFs addition, where spots correspond to permeabilized vesicles. Significant reduction was observed for GF (p=0.0120), SB80 (p=0.0140) and SB90 (p=0.0113) indicating a stabilization of vesicular compartments, and a significant increase for VX (p=0.0019). (F) A significant reduction of endocytic uptake was observed by FITC-dextran uptake for GF (p=0.0039), SB80 (p=0.0066), and VX(p=0.0111), with Enza and SB90 trending towards a reduction. (F) Measuring the FITC-dextran in acidified compartments, a strong significant increase was observed for Enza (p<0.001) and GF (p<0.001). Bar charts show mean ± SD, *p<0.05, **p<0.005, ***p<0.001. Statistical testing was performed using one-way ANOVA with Dunnett’s T3 post hoc test for multiple comparisons to a control group.

As lysosome-related pathways are in a state of flux, various effects can alter the abundance of lysosome-related compartments, though most commonly this is from lysosomal biogenesis or reduced turnover of lysosomes (Sardiello et al., 2009). We stained inhibitor-treated cells for transcription factor EB (TFEB), the master transcription factor for lysosomal biogenesis, and assessed its activation in the form of nuclear translocation (Figure 6C). Following inhibitor treatment, no significant increase in the ratio of nuclear to cytoplasmic TFEB was observed (Figure 6D). The lack of TFEB nuclear translocation indicates the increase in acidified compartments is not likely to be caused by increased lysosomal biogenesis.

We then sought to investigate the flux of the endo-lysosomal pathway by fluorescein isothiocyanate (FITC) conjugated dextran uptake and pH-dependent quenching. Vesicular pH can be assessed by detection of internalized FITC-dextran at different wavelengths, one at 530 nm, which is pH-sensitive, and the other at 610 nm which is less pH-sensitive(Eriksson et al., 2017). Similarly, we measured internalized FITC-dextran in basic and acidified compartments (Figure 6F-G). We found internalization and endosomal load to be significantly lowered for GF, SB80, and VX compared to DMSO control, but no significant difference between the inhibitor-treated samples. On the contrary, we observed a highly significant increase in FITC-dextran in acidified compartments for cells treated with Enzastaurin and GF, indicating a build-up of internalized material in acidic compartments (Figure 6G).

To investigate the effects on the internalization of α-syn, we employed a cellular model for vesicular permeabilization, using galectin3-GFP (Gal3-GFP). This model develops puncta where vesicles are permeabilized as Gal3-GFP accumulates inside the vesicles. The normalized area of broken vesicles to the total cell area shows a significant reduction in permeabilization events for GF, SB80, and SB90. In line with previous results of aggregate load, VX deviated from other inhibitors by displaying a significant increase in permeabilized vesicles following PFFs addition (p=0.0019) (Figure 6E).

Taken together, the results suggest that PKC and p38 MAPK inhibition can impact seeded α-syn aggregation, likely by preventing vesicular permeabilization and cytosolic entry.

## Discussion

Here we developed a sensitive and robust cellular reporter for FRET-based detection of α-syn aggregation. Using our cellular reporter, we performed an HTS of small molecule kinase inhibitors, identifying three candidate compounds that diminish seeded α-syn aggregation by inhibiting two actionable therapeutic pathways.

### Development of cellular α-syn aggregation reporters

When studying protein aggregation, it is essential to select the right model system based on the inherent benefits and limitations. One main factor is the model system’s complexity, from cell-free assays to animal models or patient samples. Cell-free studies have been used to explore different aspects of aggregation, including kinetics and structure (Kostka et al., 2008; Kurnik et al., 2018; Sweers et al., 2012). Yeast models leverage increased biological relevance while retaining high throughput. Therefore screening experiments for the study of α-syn and its aggregation have frequently been carried out using yeast models and have identified modifiers of α-syn toxicity, interaction partners, and protein aggregation (Cooper et al., 2006; Khurana et al., 2017; Newby et al., 2017).

We sought to establish a model system that would capture important insights into the biology of synucleinopathies. For this purpose, eukaryotic cellular models offered a midpoint between biological relevance and sample throughput. Several cell-based models are available and have been used to study native state interactions (Chung et al., 2017), such as cell-wide aggregate load (Newby et al., 2017) and fluorescent fusion constructs to study the cellular aggregation state (Prusiner et al., 2015; Tetzlaff et al., 2008; Yamasaki et al., 2019). However, no model offers direct quantification of aggregates in a workflow easily adaptable to HTS. The primary attributes needed for a model to be fit for HTS is robustness and scalability. We opted to use a FRET-based system as previously reported by Diamond and colleagues for both α-syn and tau (Holmes et al., 2014; Yamasaki et al., 2019). Detection of aggregation through FRET directly measures the aggregation state, which is highly sensitive and can be performed with a flow cytometric workflow for data analyses, as shown in our kinase inhibitor screen.

### Utilizing FRET HTS for compound discovery

With advances in automation, HTS can be carried out with thousands of samples, facilitating rapid discovery of potential therapeutic agents and molecular mechanisms. We selected a small molecule kinase inhibitor library for our screen based on the previous implication of kinases in synucleinopathies (Klegeris et al., 2008; Lee et al., 2004; Mbefo et al., 2010; Orenstein et al., 2013; Wilms et al., 2009). Among the 81 compounds screened we identified 3 potent inhibitors of seeded α-syn aggregation.

While FRET-based detection offers a unique solution for screening for which this monoclonal reporter line has been developed where it provides a simple experimental setup and analysis workflow, it also has its limitations. Among these is the reliance on both donor and acceptor being present in the same cells in the same ratios and the challenges that arise from dual transduction. The approach we apply here would not be as reliable in primary neurons or animal models, as both parts of the FRET pair are required for functionality. Low viral load risks transduced cells only receiving one FRET component, while high viral load may cause unwanted toxicity. These limitations are however overcome in a screening paradigm as presented here due to the possibility of establishing a monoclonal reporter system. To overcome the limitations related to differential transduction and expression multicistronic vectors or trans-splicing may offer viable solutions. However, attaining accurate stoichiometric control is difficult with the potential exception of 2A self-cleavable peptides (Davidsson et al., 2018).

With these limitations in mind, we were able to successfully identify 3 compounds with robust modulation of induced aggregation, clustering on p38 MAPK and PKC, which is supported by previous studies having suggested these pathways as potential therapeutic targets (Gui et al., 2020; Iba et al., 2020; Iwata et al., 2001; Obergasteiger et al., 2018), (Do Van et al., 2016; Weinreb et al., 2005; Youdim et al., 2003). While evidence for a protective role of modulation of these pathways has been suggested, the mechanism by which they might cause the protection is currently not known.

### The implication of the endo-lysosomal system

Identifying the molecular changes responsible for the protective effect is difficult when working with broad pathways like p38 MAPK and PKC. Therefore, the use of mass-spectroscopy for an unbiased investigation of the cellular proteome is a valuable methodology for target identification.

The phospho-proteome showed statistically significant alterations in the phosphorylation state of proteins related to protein synthesis, compartment organization, and biogenesis, in line with the inhibitors’ known targets. Both PKC and p38 MAPK are known upstream modulators for a series of molecular functions. PKC is a major regulator for cell division and proliferation (Reyland, 2009), while p38 mediates apoptotic/survival gene programs (Yue & López, 2020), differentiation (Neganova et al., 2017) as well as being involved in inflammation (He et al., 2018).

The proteome instead revealed pathways with potential implications for the observed effect of the inhibitors. A set of mitochondria and metabolism-related proteins mapped to KEGG’s PD, AD and HD pathways. The involvement of mitochondria and reactive oxygen species for these diseases is well known and documented (Angelova & Abramov, 2018). However, the downregulation of these KEGG signatures was only present for three of four inhibitors with the protective effects while being absent for SB80 and VX. Instead, an increase in proteins mapped to KEGG’s lysosome pathway was recaptured by all four protective inhibitors. The importance of lysosomal function in relation to synucleinopathies is well supported, with familial variants in glucocerebrosidase (GBA), a lysosomal hydroxylase, being one of the largest risk factors for developing PD (Beavan & Schapira, 2013; Brockmann & Berg, 2014; Velayati et al., 2010).

When assessing the abundance of acidic compartments, significant increases were only observed for Enza, GF and SB90, whereas no treatment led to identifiable activation of TFEB driven lysosomal biogenesis. This is contrary to other reports which indicate there is TFEB translocation upon treatment with SB90, corresponding to classical TFEB activation and lysosomal biogenesis (Yang et al., 2020).

Whether the increased abundance of lysosomes is beneficial or detrimental is still under debate. It is well established that α-syn degradation can be facilitated through chaperone-mediated autophagy, which is lysosome-dependent (Beavan & Schapira, 2013). However, tau seeding in similar models as ours identifies acidified compartments as a necessary component for tau seeds to enter cells and induce pathology. Disruption of these acidified compartments protected against induced tau aggregation (Polanco et al., 2021).

We further performed a galectin-3 vesicular permeabilization assay and found all 4 protective inhibitors to prevent vesicle permeabilization, with one compound, VX, instead trending towards an exacerbation. The implication of altered vesicle permeabilization would indicate the inhibitors’ protective effects on induced aggregation occurs at or before the point of vesicle permeabilization. This prevention of vesicular disruption could well be caused by increased degradative capacity of lysosomes or sequestration of exogenous detrimental aggregates in vesicular compartments as indicated by the FITC-dextran uptake for p38 MAPK and PKC respectively (Figure 7).

**Figure 7.**
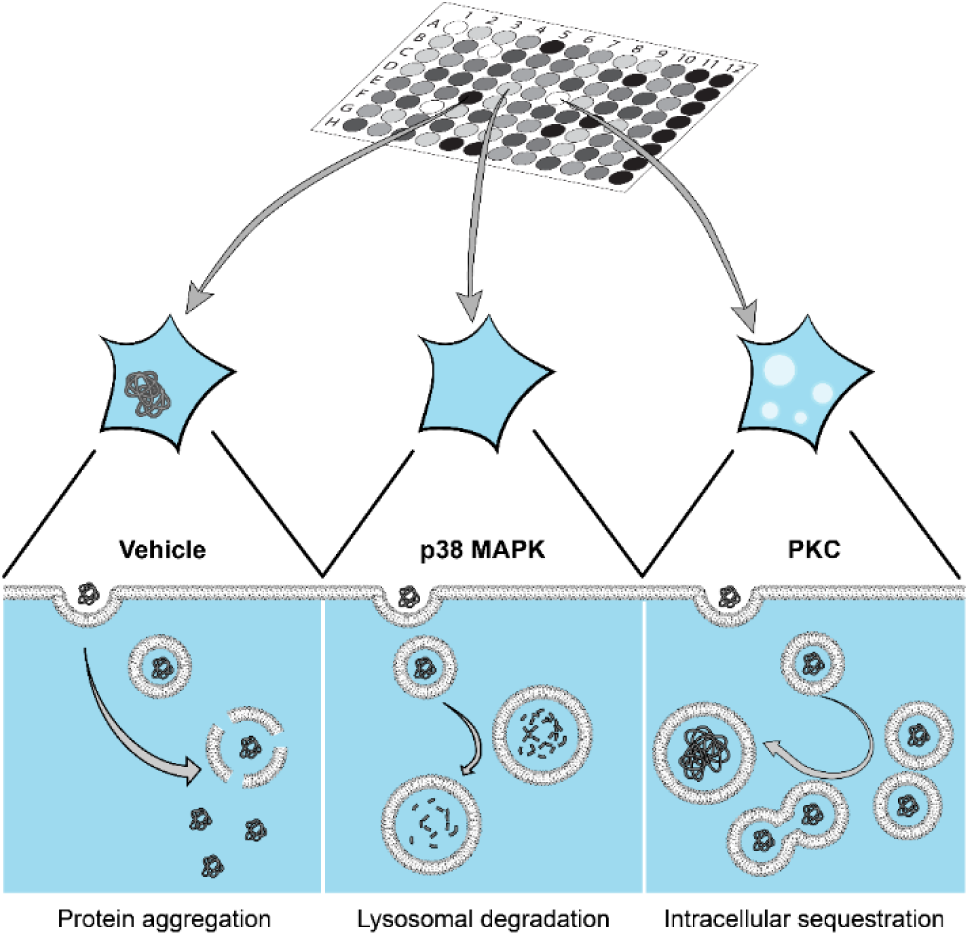
Proposed model for the protective effects of p38 MAPK- and PKC-inhibition on induced α-syn aggregation. The PFF seeded α-syn aggregation paradigm, relies on the extracellular addition of seed material, capable of gaining cellular entry and recruiting endogenous α-syn thereby initiating intracellular aggregation. Upon inhibition of p38 MAPK and PKC by SB80, SB90 and GF, vesicular disruption is prevented. Extracellularly added α-syn may be either degraded or sequestered upon endocytosis, in both cases preventing induction of α-syn aggregation.

## Conclusion

In conclusion, our novel HTS system based on the NPR-H-001 reporter cells provides a new method to study α-syn aggregation and modulation thereof by compounds or other perturbations. Unbiased screening of effectors of α-syn aggregation, as shown here, further allows the identification of molecular mechanisms involved in α-syn pathology. From the screening, we identified three potent inhibitors of induced α-syn aggregation, and through their targets could confirm p38 MAPK and PKC as potential therapeutic targets. Target discovery from screens such as this serves to further deepen our understanding of the molecular mechanisms. In this case, we found further support for the importance of the endo-lysosomal systems in α-syn pathology. Further, more comprehensive screens may serve to broaden our understanding of the molecular mechanisms essential for the induction and maintenance of α-syn pathology.

## Acknowledgements

We thank the Center for Translational Proteomics, CTP, at Medical Faculty and Region Skåne, Lund University. We thank Dr. Anna Hammarberg for her excellent technical support. We also thank Andreas Heuer and Marcus Davidsson for scientific input and feedback throughout the process, and Luis Quintino for guidance on the topic of lysosomes. We acknowledge financial supports by The Swedish Research Council (K2015-61X-22297-03-4; 2019-01551), EU-JPND research (aSynProtec and REfrAME) and EU-Horizon2020 (MSCA-ITN-2016, SynDeGen), ParkinsonFonden, the Strategic Research Area Multipark (Multidisciplinary research in Parkinson’s disease at Lund University) and the National Natural Science Foundation of China (81430025, 31800898, and U1801681) and the Key Field Research Development Program of Guangdong Province (2018B030337001).

## Materials and Methods

### α-syn production

Recombinant α-syn was produced as previously described (Gaspar et al., 2018). Briefly, codon optimized human α-syn was expressed in *E. Coli* using a Pet3a inducible plasmid. Bacterial cultures were maintained at 37°C with 125 rpm shaking until an OD_600_ value between 0.6 and 1.00 was obtained, at which point α-syn expression was induced by 0.4 mM IPTG for 4 hours.

Cultures were harvested by centrifugation and α-syn purified by heat treatment ion exchange chromatography. Finally, to ensure monomeric α-syn was produced, the isolated protein was run on a size exclusion column (Superdex 75, GE Healthcare) with a Tris buffer (10 mM Tris, 150 mM, NaCl pH 7.5). From the resulting fractions, the central fraction of the monomer peak was collected. Yield was determined by absorbance at 280 nm (Nanodrop, Thermo fisher) and calculated using an extinction coefficient ε = 5800.1 M^-1^cm^-1^.

### Fibril production

Freshly prepared human recombinant α-syn was adjusted to a concentration of 0.5 µg/µl with Tris buffer (10 mM Tris, 150 mM, NaCl pH 7.5) and incubated for 14 days at 37 °C with 1000 rpm agitation using 3mm magnets in low binding 1.5 ml Eppendorf tubes. Fibrillation of the monomeric α-syn was confirmed with Thioflavin T readings and transmission election microscopy.

For all experiments using PFFs, cup-horn sonication was performed immediately prior to addition using a Qsonica Q125 Sonicator (Qsonica, Newtown, CT) for 3 min in cycles of 1 sec on, 1 sec off at 70% amplitude.

### Cloning

Using constructs previously generated in our laboratory carrying α-syn^A53T^-CFP and α-syn^A53T^- YFP as donor plasmids fluorescently tagged α-syn was PCR amplified and overhangs added for XbaI/XhoI digestion. By enzymatic digestion and gel purification of the PCR product, the amplicon was inserted into a linearized pHsCXW backbone with the same overhangs. The resulting plasmids (pHsCXW: α-syn^A53T^-CFP and pHsCXW: α-syn^A53T^-YFP) gave a strong uniform CMV-driven expression α-syn carrying the fluorophores of interest.

### Lentivirus production

Lentivirus production was carried out in HEK293-T cells, using packaging plasmids pMD2G, pMDL and pRsvREV, and the lentiviral transfer plasmid carrying the expression cassette of interest. Cells were seeded in a Nunc T175 flask at a density of 14·10^6^ cells/flask and incubated over-night. The following day, 2 hours before transfection the culture media was replaced for 16.2 ml fresh media. For lentivirus assembly, 5,1 µg pMD2G, 7.1 µg pMDL, 4.0 µg pRsvREV and 18.0 µg transfer plasmid was mixed in 1 ml of DMEM without supplements. 102.6 µl PEI was added to the reaction mix and total volume adjusted to 1.8 ml before vortexing thoroughly. The transfection mix was incubated for 15 min at room temperature before gently adding to the cells. Media was harvested at 48 and 72 hours and replaced by new media. After the second harvest, collected media was spun at 800g, 4 °C for 10 min to pellet cell debris and filtered through a 0.22 um filter. The filtrate was transferred to a Beckman Coulter ultracentrifuge tube and spun at 25000 rpm in SW-32 swing-buckets to concentrate the virus particles. The supernatant was discarded, and the tubes were air dried before adding 80 µl sterile PBS to each tube to resuspend the pellet over-night. After resuspension, the virus suspension was aliquoted and tittered.

### Cell culture

All cells were cultured at 37 °C, 95% relative humidity and 5% CO_2_ in DMEM supplemented with 10% FBS and 1% penicillin/streptomycin. Maintenance of the cultures was done by sub-culturing at 80-90% confluence by gentile detachment of the cells, transferring to new culture flasks with fresh media. 96 well multi-well plates were coated with Collagen G for 1 hour and washed twice with PBS before seeding cells. All experiments for direct addition of α-syn fibrils were seeded with 30.000 cells/cm^2^ and experiments using lipofectamine for fibril delivery were seeded with 90.000 cells/cm^2^.

### Generation of FRET based cell model for α-synuclein aggregation

Lentiviral transduction with 1 MoI of each α-syn^A53T^-CFP and α-syn^A53T^-YFP was used to facilitate stable integration of the FRET reporter molecules in Hek293T cells. After 1 week of maintaining the culture, cells were harvested by trypsin, pelleted, resuspended in PBS supplemented with 2 % FBS before being passed through a 100 um cell strainer (Falcon). The resulting single cell suspension was stained with Draq7 (1:10.000) for viability assessment and sorted on a FACSAria II as single cells into a 96 well culture plate with 50 % fresh media and 50 % conditioned media (Conditioned media was prepared taking cells from the parental HEK293T culture, spinning at 800g for 10 min and filtering through a 0.22 um syringe filter). Sorted cells were expanded and frozen as monoclonal reporter cells.

### Immunocytochemistry

Cells were washed with PBS, fixed with 4% paraformaldehyde and blocked with PBS+1% BSA+0.1% Tween for 1 hour. Cells were incubated at 4 deg C over night with primary antibody (Supplemental Table 2) in blocking buffer. Excess primary antibody was removed by washing three times with PBS and stained by species-specific secondary antibody for 1 hour and DAPI (1µg/ml) at room temperature. Prior to imaging cells were washed three times with PBS. For staining with CongoRed cells were incubated for 15 minutes with 1 µM CongoRed (Sigma-Aldrich) in 60% PBS/39% ethanol/20mM NaOH. Cells were rinsed three times in staining buffer without CongoRed followed by five times with PBS before imaging.

### Protein extraction and Immunoblotting

For extraction of detergent-soluble and -insoluble material was performed as previously described (Fares et al., 2016), cells were harvested in cold lysis buffer (20 mM Trizma base, 150 mM NaCl, 1mM EDTA, 0.25% NP-40, 0.25% Triton X-100, pH 7.4) with protease and phosphatase inhibitors. After 20 min incubation on ice, lysate was spun for 20 min at 4 deg C, 14000g. Supernatant was stored as detergent soluble fraction, while the pellet was resuspended in lysis buffer with added 5% (wt/vol) SDS and sonicated by cup-horn at 60% amplitude, 3 sec on/off cycles for 15 sec.

For immunoblotting, even protein amounts were loaded and separated on mini-PROTEAN TGX precast 4-15% Bis-Tris gels (#4568085, Bio-Rad, Copenhagen, Denmark). Using a Bio-Rad Trans-Blot semidry transfer system proteins were transferred to a nitrocellulose membrane (#1704159, Bio-Rad) and blocked for 1 hour (PBS, 0.1% Tween20, 3% BSA) at room temperature. Primary antibody (supplementary table 2) staining was performed in blocking buffer at 4 deg C over night on a shaking table. After three washes with (PBS, 0.1% Tween20) incubation with secondary species specific HRP-conjugated antibody was performed in blocking buffer for 1 hour prior to development on a Bio-Rad ChemiDocMP imaging system.

### Lysotracker assessment of acidified compartments

To assess the extent of acidic compartments within cells, Lysotracker Deep Red (#L12492, ThermoFisher) was added to culture medium 30 prior to analysis (1:1000). For live-cell imaging Hoechst 33342 (#62249, ThermoFisher) was added at a concentration of 0.1 µg/ml together with Lysotracker, while imaging was performed on a Nikon TI microscope equipped with a OKOlab live cell chamber. Flow cytometric analysis was performed by harvesting cells by trypsin detachment, resuspension into PBS supplemented with 1 µg/ml DAPI for staining dead cells.

### FITC-dextran uptake assay

To follow endo-lysosomal uptake we added 10kDa FITC-dextran (#FD10S-100MG, Sigma-Aldrich) at 0.1 mg/ml and incubate at 37 deg C. After 1 hour of incubation, cells were washed twice with PBS and harvested by trypsin digest and resuspended in PBS. Cells in suspension were analyzed by flow cytometry on a LSRFortessa ex. 488, em. 530 and ex. 610, for assessment of the pH sensitive and pH insensitive parts of the FITC spectrum(Eriksson et al., 2017).

### Small molecule compound library screen

An established small molecule kinase inhibitor library (BML-2832, Enzo Life sciences) was selected for the screen. All compounds included in the library as well as the resulting z-scores can be found in supplementary table 1. All compounds were dissolved in DMSO at a concentration of 10 mM. For each screen, 10.000 cells/well were seeded in a collagen G coated 96 well plate the day prior to addition of inhibitors and α-syn fibrils. The following day inhibitors was added by pipetting robot to each well for the final concentration of 2, 4 and 10 µM after addition of α-syn. Half an hour after addition of inhibitors, α-syn fibrils were added and observed for 48 hours by live-cell imaging. At endpoint, the cells were harvested with trypsin and fixed in suspension for 20 min on ice. Fixed cells were washed twice in PBS before analysis of normalized FRET intensity (% FRET^+^×FRET MFI) by flow cytometry (LSRFortessa). Compounds were ranked based on calculated 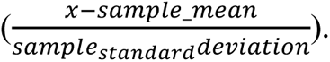 Threshold for hit compounds was set at z-score=±1.5.

### Image analysis

Image analysis was performed using Python 3.1 with image analysis pipelines made available through GitHub. Co-localization analysis was performed by calculating the PCC for every pixel in images between the two staining’s in question (https://github.com/AlexanderSvan/PCC-colocalization-for-images). Aggregate detection was performed by application of gaussian blur and a white top hat filter, prior to setting a threshold for aggregates. Object detection was performed using the package Sci-kit image, and aggregates were filtered based on aggregate size (https://github.com/AlexanderSvan/Live-cell-aggregate-count). For TFEB nuclear translocation cells were identified and subdivided into cytosol and nucleus based on DAPI staining. Translocation was calculated as the ratio of TFEB intensity between nucleus/cytosol (https://github.com/AlexanderSvan/TFEB-translocation).

### Proteomics MS

#### Samples

18 cell pellets in total: 5 different drugs treatments and 1 control conditions, 3 biological replicates for each

#### Sample preparation

Unless stated otherwise, all the chemicals and solvents were purchased from Sigma-Aldrich, Steinheim, Germany.

Suspension traps (S-Traps) based digestion protocol was used for the sample processing according to the manufacturer’s instructions.(Zougman et al., 2014) Cell pellets were lysed by using a detergent based lysis buffer (25 mM dithiothreitol (DTT), 5% sodium dodecyl sulfate (SDS) in 50 mM triethylammonium bicarbonate (TEAB, Thermo Fisher Scientific, Rockford, IL), pH=8.0). The lysates were boiled for 5 min at 95°C in a thermomixer with the mixing frequency of 500 rpm and then sonicated to denature proteins, shear DNA and enhance cell disruption by using a water-bath (4°C) sonicator Bioruptor (Diagenode) at High Power for 40 min (sonication cycle: 15 sec ON, 15 sec OFF). The samples were centrifuged at 16,000 g for 10 min at room temperature and the supernatant were transferred to new tubes. The concentration of protein was determined by using Pierce™ 660nm Protein Assay Reagent with Ionic Detergent Compatibility Reagent (IDCR) (Thermo Fisher Scientific, Rockford, IL). Around 300 µg of cell lysates for each sample were alkylated with 50 mM iodoacetamide (IAA) for 30 min in the dark. Afterward, each sample was acidified by aqueous phosphoric acid (Merck) with the final concentration of 1.2% and then seven volumes of S-Trap binding buffer (90% methanol, 100 mM TEAB, pH=7.1) was added. After gentle mixing, the protein solution was loaded directly onto the S-Trap spin column (Protifi, Huntington, NY) without any column pre-equilibration, and spun at 4000g for 30 seconds. The S-Trap column was washed 3 times by using 400 µL S-Trap binding buffer. Each sample was digested with 95 µL 50 mM TEAB containing Lys-C (FUJIFILM Wako Chemicals U.S.A. Corporation) at an enzyme: protein ratio of 1:50 w/w for 2 hours at 37°C and further digested with 30 µL 50 mM TEAB containing trypsin (Sequencing Grade Modified, Promega, Madison, WI) at a trypsin: protein ratio of 1:50 w/w overnight at 37°C. The peptides were eluted by three stepwise buffers with 80 µL of each, including 50 mM TEAB, 0.2% formic acid (FA) in water and 0.2% FA in 50% acetonitrile (ACN). The three steps elution’s were pooled together and dried in a SpeedVac (Concentrator plus Eppendorf).

The peptide concentration was determined by using the Pierce Quantitative Colorimetric Peptide Assay (Thermo Fisher Scientific, Rockford, IL) and 180 µg peptides from each sample were loaded onto MacroSpin column (NestGroup, Southborough, MA) for desalting. In brief, the spin column was primed and equilibrated with 80% ACN/0.1% TFA and 0.1% TFA, respectively and was washed with 0.1% TFA after sample loading. The peptides were eluted with 80% ACN/0.1% TFA, dried in the SpeedVac and stored in -80°C freezer until further process.

Phosphor enrichment was performed on the Agilent AssayMAP Bravo Platform (Agilent Technologies, Inc) and phosphorylated peptides were enriched by using 5 μL Fe(III)-NTA cartridges. Following the Phospho Enrichment v2.0 protocol, the cartridges were primed with 50% ACN/0.1% TFA and equilibrated with 80% ACN/0.1% TFA. Peptides from desalting step were re-suspended with 80% ACN/0.1% TFA and loaded onto the cartridge for the enrichment. The phosphorylated peptides were eluted with 5% ammonia directly into 50% FA after being washed with 80% ACN/0.1% TFA, lyophilized in the SpeedVac and stored in -80°C freezer until LC-MS/MS analysis.

#### LC MS/MS analysis

The LC MS/MS analysis was performed on Tribrid mass spectrometer (MS) Orbitrap Fusion equipped with a Nanospray source and coupled with an EASY-nLC 1000 ultrahigh pressure liquid chromatography (UHPLC) pump (Thermo Fischer Scientific, San Jose, CA). One microgram peptides for global proteomics or all the enriched phosphorylated peptides were loaded and concentrated on an Acclaim PepMap 100 C18 precolumn (75 μm x 2 cm, Thermo Scientific, Waltham, MA) and then separated on an Acclaim PepMap RSLC column (75 μm x 25 cm, nanoViper, C18, 2 μm, 100 Å) with the column temperature of 45°C. Peptides were eluted by a nonlinear 2h gradient at the flow rate of 300 nL/min from 2% solvent B (0.1% FA/ACN)/98% Solvent A (0.1% FA in water) to 40% solvent B.

The Orbitrap Fusion was operated in the positive data-dependent acquisition (DDA) mode for both global proteomics and phosphor-proteomics. Full MS survey scans from m/z 350-1500 with a resolution 120,000 were performed in the Orbitrap detector. The automatic gain control (AGC) target was set to 4 × 10^5^ with an injection time of 50 ms. The most intense ions (up to 20) with charge state 2-7 from the full MS scan were selected for fragmentation. MS2 precursors were isolated with a quadrupole mass filter set to a width of 1.2 m/z for global proteomics and 0.7 for phosphor-proteomics. Precursors were fragmented by Higher Energy Collision Dissociation (HCD) and detected in Orbitrap detector with the resolution of 30,000 for global proteomics and 60,000 for phosphor-proteomics. The normalized collision energy (NCE) in HCD cell was set 30%. The values for the AGC target and injection time were 5 × 10^4^ and 54 ms for global proteomics and 110 ms for phosphor-proteomics, respectively. The duration of dynamic exclusion was set 45 s and the mass tolerance window 10 ppm.

#### Data analysis

All the MS raw files were submitted to Proteome Discoverer 2.3 (Thermo Scientific) for identification and quantification. The search was performed against the Homo sapiens UniProt revised database with the Sequest HT search engine and decoy database containing reversed version of all protein sequences was used to monitor false discovery rate (FDR). Carbamidomethylation of cysteine residues was set as fixed modification and methionine oxidation, protein N-terminal acetylation and phosphorylation of serine, threonine and tyrosine were set as variable modifications. For peptide identification, precursor mass tolerance was set 15 ppm and fragment mass tolerance 0.05 Da respectively. A maximum of two missed cleavage sites was allowed. The IMP-ptmRS algorithm was used to score phosphorylation sites with a site probability threshold >75. After database searching, 1% false discovery rate (FDR) for both peptide-spectrum match (PSMs) and peptides was applied to filter out the wrong peptides. For statistical analyses, label-free quantification (LFQ) data was log2-transformed and normalized to median value of each sample. Multiple-sample test was performed within the Perseus software, with Dunnett’s T3 post-hoc test.

### Functional analysis

We performed functional analysis using DAVID (Huang et al., 2009b, 2009a) using the differentially expressed genes from analysis of the proteome and phosphor-proteome

---

**Supplementary Figure 1:**
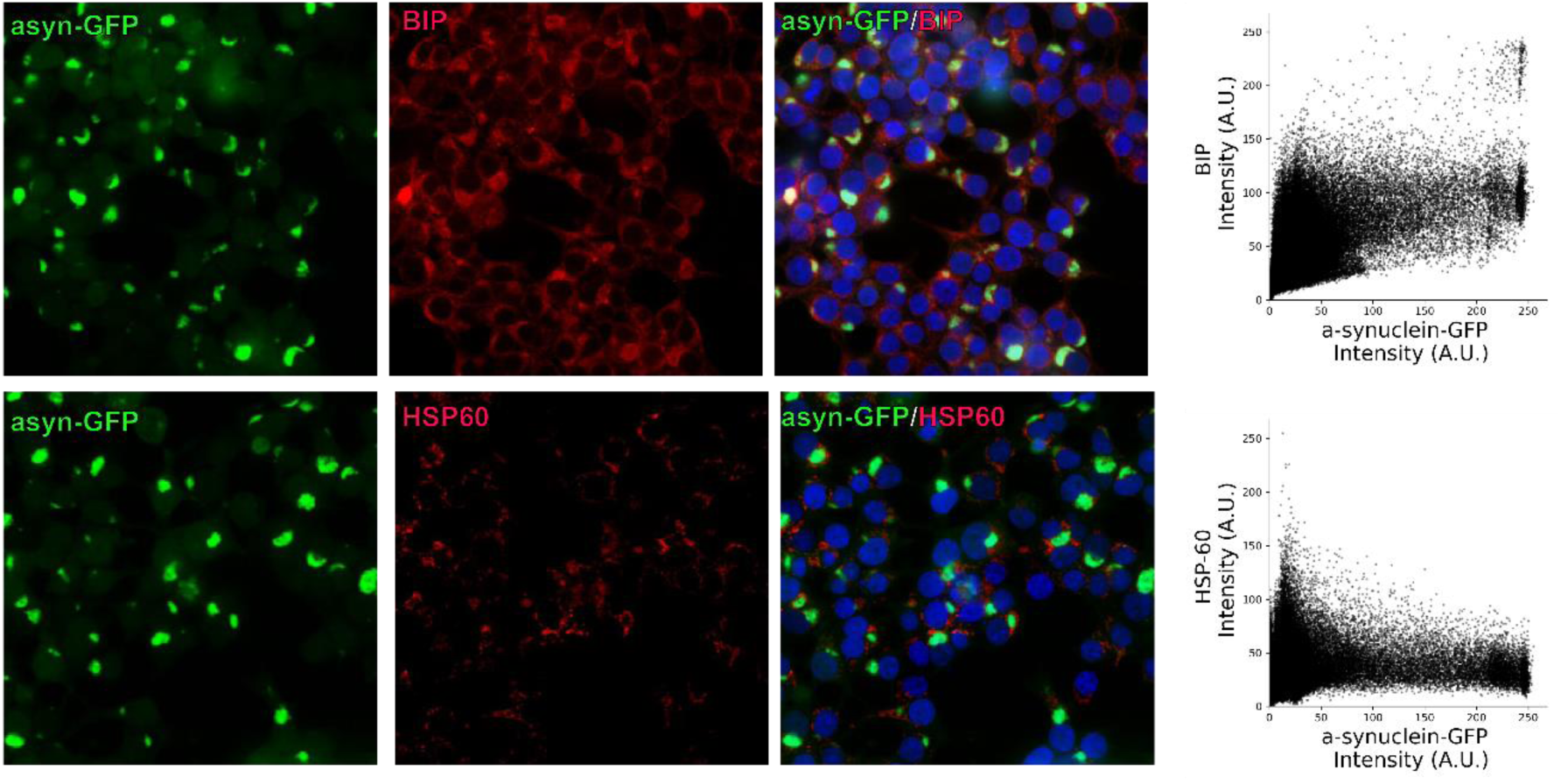
Co-localzation of α-syn to molecular chaperones. (A-B) Induced aggregates co-localize to areas of increased BIP staining, leading to a positive PCC of 0.6±0.1 (C-D) HSP60 is seen concentrated in small foci, and primarily observed surrounding GFP positive inclusions, also reflected in the lower PCC of 0.29±0.008. All PCC measurements are given in mean±SD.

**Supplementary table 1.**

| **Name** | **Compound description** | **Z-score** |
| --- | --- | --- |
| *Negative control* | Nan | -1.88536 |
| *U-0126* | MEK | -1.88513 |
| *Negative control* | Nan | -1.88395 |
| *Negative control* | Nan | -1.88236 |
| *Negative control* | Nan | -1.86253 |
| *SB-202190* | p38 MAPK | -1.75475 |
| *GF 109203X* | PKC | -1.54164 |
| *SB-203580* | p38 MAPK | -1.53681 |
| *Staurosporine* | Pan-specific | -1.36907 |
| *Apigenin* | CK-II | -1.30894 |
| *Ro 31-8220 mesylate* | PKC | -1.16857 |
| *PKC-412* | PKC | -1.15901 |
| *Kenpaullone* | GSK-3beta | -1.08908 |
| *TYRPHOSTIN 9* | PDGFRK | -1.05809 |
| *ML-9 HCl* | MLCK | -0.97196 |
| *RG-1462* | EGFRK | -0.79068 |
| *Indirubin-3'-monooxime* | GSK-3beta | -0.63862 |
| *Hydroxy-2-naphthalenylmethylphosphonic acid* | IRK | -0.61666 |
| *D-erythro-Sphingosine* | PKC | -0.5356 |
| *TYRPHOSTIN 23* | EGFRK | -0.49958 |
| *TYRPHOSTIN 47* | EGFRK | -0.49252 |
| *Piceatannol* | Syk | -0.46871 |
| *BML-259* | Cdk5/p25 | -0.46475 |
| *TYRPHOSTIN 51* | EGFRK | -0.42424 |
| *Indirubin* | GSK-3beta, CDK5 | -0.42325 |
| *2-Hydroxy-5-(2,5-dihydroxybenzylamino)benzoic acid* | EGFRK, CaMK II | -0.40071 |
| *Triciribine* | Akt | -0.39062 |
| *Palmitoyl-DL-carnitine* | PKC | -0.36036 |
| *TYRPHOSTIN AG 1288* | Tyrosine kinases | -0.35195 |
| *GW 5074* | cRAF | -0.34121 |
| *Wortmannin* | PI 3-K | -0.33565 |
| *TYRPHOSTIN 1* | Negative control for tyrosine<br>kinase inhibitors. | -0.28111 |
| *SP 600125* | JNK | -0.26322 |
| *SU 4312* | Flk1 | -0.26231 |
| *Lavendustin A* | EGFRK | -0.2559 |
| *2-Aminopurine* | p58 PITSLRE beta1 | -0.25187 |
| *H-9 HCl* | PKA, PKG, MLCK, and PKC. | -0.24961 |
| *TYRPHOSTIN AG 1295* | Tyrosine kinases | -0.236 |
| *AG-494* | EGFRK, PDGFRK | -0.21098 |
| *Hypericin* | PKC | -0.20756 |
| *H-8* | PKA, PKG | -0.19294 |
| *Quercetin 2H2O* | PI 3-K | -0.18958 |
| *Roscovitine* | CDK | -0.16391 |
| *Olomoucine* | CDK | -0.14604 |
| *H-7 2HCl* | PKA, PKG, MLCK, and PKC. | -0.13913 |
| *SC-514* | IKK2 | -0.13596 |
| *AG-1296* | PDGFRK | -0.1359 |
| *ZM 336372* | cRAF | -0.13218 |
| *Y-27632 2HCl* | ROCK | -0.12655 |
| *Daidzein* | Negative control for Genistein. | -0.12489 |
| *2,2',3,3',4,4'-Hexahydroxy-1,1'-biphenyl-6,6'-dimethanol dimethyl ether* | PKC alpha, PKC gamma | -0.09088 |
| *ZM 449829* | JAK-3 | -0.08391 |
| *Terreic acid* | BTK | -0.04542 |
| *N9-isopropyl-olomoucine* | CDK | 0.012188 |
| *ML-7 HCl* | MLCK | 0.032992 |
| *PD-98059* | MEK | 0.05293 |
| *SU1498* | Flk1 | 0.05404 |
| *AG-825* | HER1-2 | 0.054351 |
| *H-89 2HCl* | PKA | 0.204342 |
| *Iso-olomoucine* | Negative control for olomoucine. | 0.206719 |
| *AG-370* | PDGFRK | 0.236599 |
| *BAY 11-7082* | IKK pathway | 0.265294 |
| *HA-1004 2HCl* | PKA, PKG | 0.281961 |
| *BML-257* | Akt | 0.296339 |
| *5,6-dichloro-1-β-D-ribofuranosylbenzimidazole* | CK-II | 0.305907 |
| *AG-126* | IRAK | 0.360894 |
| *TYRPHOSTIN 46* | EGFRK, PDGFRK | 0.391667 |
| *LFM-A13* | BTK | 0.410376 |
| *TYRPHOSTIN AG 1478* | EGFRK | 0.45931 |
| *HA-1077 2HCl* | PKA, PKG | 0.474783 |
| *TYRPHOSTIN 25* | EGFRK | 0.519171 |
| *AG-879* | NGFRK | 0.594178 |
| *Positive control* | Nan | 0.692666 |
| *Positive control* | Nan | 0.787487 |
| *BML-265* | EGFRK | 0.858547 |
| *PP2* | Src family | 0.910366 |
| *PP1* | Src family | 0.914834 |
| *Positive control* | Nan | 0.94276 |
| *AG-490* | JAK-2 | 1.03706 |
| *Positive control* | Nan | 1.097282 |
| *KN-93* | CaMK II | 1.104091 |
| *Positive control* | Nan | 1.138769 |
| *Genistein* | Tyrosine Kinases | 1.261532 |
| *Rottlerin* | PKC delta | 1.264283 |
| *Positive control* | Nan | 1.380756 |
| *Positive control* | Nan | 1.422028 |
| *Positive control* | Nan | 1.481894 |
| *KN-62* | CaMK II | 1.962847 |
| *LY 294002* | PI 3-K | 2.002057 |
| *Erbstatin analog* | EGFRK | 2.347802 |
| *5-Iodotubericidin* | ERK2, adenosine kinase, CK1,<br>CK2, | 2.595458 |
| *Rapamycin* | mTOR | 3.901389 |

| Antibody | Supplier | Species | Dilution |
| --- | --- | --- | --- |
| **asyn 211** | Santa Cruz (#SC12767) | Mouse | 1:1000 |
| **Pasyn** | Abcam (#AB51253) | Rabbit | 1:1000 |
| **HSP60** | Cell Signaling (#4870) | Rabbit | 1:1000 |
| **BIP** | Abcam (#ab21685) | Rabbit | 1:1000 |
| **TFEB** | Thermo Fisher (#A303-673A) | Rabbit | 1:1000 |

## Notes

### Competing Interest Statement

The authors have declared no competing interest.

